# Mammalian glial protrusion transcriptomes predict localization of *Drosophila* glial transcripts required for synaptic plasticity

**DOI:** 10.1101/2022.11.30.518536

**Authors:** Dalia S. Gala, Jeffrey Y. Lee, Maria Kiourlappou, Joshua S. Titlow, Rita O. Teodoro, Ilan Davis

## Abstract

The polarization of cells often involves the transport of specific mRNAs and their localized translation in distal projections. Neurons and glia both contain long cytoplasmic processes with important functions. mRNA localization has been studied extensively in neurons, but very little in glia, especially in intact nervous systems. Here, we predicted 1,700 localized *Drosophila* glial transcripts by extrapolating from our meta-analysis of 8 existing studies characterizing the localized transcriptomes and translatomes of synaptically-associated mammalian glia. We tested these predictions in glia of the neuromuscular junction of *Drosophila* larvae and found that localization to mammalian glia is a strong predictor of mRNA localization of the high confidence *Drosophila* homologues. We further showed that some of these localized transcripts are required in glia for plasticity of the neuromuscular junction synapses. We conclude that peripheral glial mRNA localization is a common and conserved phenomenon and propose that it is likely to be functionally important.

## 1 INTRODUCTION

It is well established that specific mRNAs are actively transported and localized to distal compartments of neurons, including those encoding cytoskeletal proteins, neurotransmitters, membrane proteins and ribosomes. In these studies, mRNA transport and local translation have been proposed as mechanisms for local regulation of synaptic plasticity, which is the process underlying memory and learning (Smith, Aakalu and Schuman, 2001; Sutton and Schuman, 2006; Wang, Martin and Zukin, 2010; Daniel E Shumer, 2017; Rangaraju, tom Dieck and Schuman, 2017; Terenzio, Schiavo and Fainzilber, 2017; Biever, Donlin-Asp and Schuman, 2019; Holt, Martin and Schuman, 2019; Wang *et al*., 2009).

Asymmetric mRNA localization is likely to be as important in glia, as it is in neurons, but has received very little attention in glia. Like neurons, glial cells can display diverse morphologies ranging from immune-cell-like microglia, to elongated myelinating oligodendrocytes and astrocytes that contact thousands of synapses. Therefore, astrocytes are thought to integrate information from distinct neuronal populations (Ransom and Ransom, 2012; Jessen, Mirsky and Lloyd, 2015; Allen and Lyons, 2018; Volterra and Meldolesi, 2005). Astrocytes also perform key roles in the uptake and release of neurotransmitters and the maintenance of ionic balance. Glia form the blood-brain barrier that regulates neuronal metabolism and local solute homeostasis, as well as secretion of gliotransmitters and mediation of signaling pathways (De Pittà, Brunel and Volterra, 2016; Wang, Fu and Ip, 2022; Ota, Zanetti and Hallock, 2013). Oligodendrocytes are thought to modulate synaptic plasticity via myelination (Fields, 2005; Fields, 2008; de Faria *et al*., 2021; Bacmeister *et al*., 2020; Pan *et al*., 2020; Fields and Bukalo, 2020). Microglia can preferentially phagocytose synaptic endings and modulate connectivity in the visual cortex (Andoh and Koyama, 2021; Graeber, 2010; Morris *et al*., 2013; Schafer *et al*., 2012; Vasek *et al*., 2021). All of these functions occur in remote cellular processes that are separated from the core gene expression machinery in the cell body. Although these varying glial morphologies and functional roles are suggestive of mRNA localization (Blanco-Urrejola *et al*., 2021; Meservey, Topkar and Fu, 2021), it is not clear whether mRNA localization is common in glia, nor whether a common set of mRNAs exists that localizes to the periphery of most of the glial sub-types or in neurons. It is also not known whether such glial mRNA localization is conserved and functionally important.

To address these questions, we have carried out a meta-analysis of, to our knowledge, all the published mammalian glial peripheral or synaptic transcriptomes and translatomes. We found several classes of mRNAs that are localized in multiple glial subtypes and determined a common set of 4801 transcripts present in the majority of the analyzed libraries, representing multiple types of mammalian glial cells. We used this cohort to find high confidence *Drosophila* homologues and filtered the list for glial expression using the Fly Cell Atlas data, a single-cell nuclear sequencing atlas derived from entire adult flies (Li *et al*., 2022). We specifically selected for expression in 3 glial subtypes known to create distal protrusions in the larval NMJ, namely perineural, subperineural and wrapping glia, which are continuous with the ensheathing glia in the PNS (Yildirim *et al*., 2019; Sepp, Schulte and Auld, 2000), resulting in 1,700 mRNAs predicted to be localized in these three glial subtypes.

To test our predictions of mRNA localization in *Drosophila* glial projections, we studied in greater detail 15 conserved transcripts, which in an unrelated survey of 200 transcripts across the nervous system, we discovered were likely to be localized at glial distal projections near the NMJ axonal synapses (Titlow *et al*., 2022). We first confirmed that these 15 transcripts were indeed localized, using definitive glial markers and more extensive, higher precision 3D microscopy. We then determined whether these experimentally determined localized transcripts are predicted as localized from our meta-analysis. We found that more than 70% (11 of the 15) conserved localized transcripts were correctly predicted as localized, through being in the list of 1,700 transcripts. Furthermore, in follow up experiments, we tested whether the localized transcripts are required in the glia for the correct plasticity of the adjacent neuronal synapses, using a spaced potassium stimulation assay (Ataman *et al*., 2008; Roche *et al*., 2002). We found that 5 of the 15 localized transcripts impaired synaptic plasticity in the neighboring wild type neurons, reducing their ability to make new synapses. Our results suggest that mRNA localization to protrusions represent an important and conserved mechanism that allows glia to rapidly regulate the plasticity of adjacent distal axonal synapses.

## 2 MATERIALS AND METHODS

### 2.1 Data meta-analysis methods

We developed a custom RNA-seq analysis pipeline to process public transcriptomic datasets and allow the integration of diverse raw datasets from 8 different studies. Raw FASTQ files were downloaded from Gene Expression Omnibus (GEO) and were filtered to remove ribosomal RNA reads using BBduk (Bushnell, Rood and Singer, 2017). Processed reads were mapped to *Mus musculus* (ENSEMBL release 96) or *Rattus norvegicus* (ENSEMBL release 98) genomes using Kallisto to obtain Transcripts per million (TPM) abundance measures (Bray *et al*., 2016). A TPM value > 10 for a transcript was considered a transcript to be present in a given compartment. In order to take into account intronic reads in libraries derived from soma, Kallisto bustools was used to create transcriptome indices containing intron sequences (Melsted *et al*., 2021). For the relative enrichment analysis comparing soma and protrusion compartments, DESeq2 was used to assess differential expression of transcripts from estimated transcript counts output (Love, Huber and Anders, 2014). The independent hypothesis weighting (IHW) method was used to correct for the multiple testing (Ignatiadis *et al*., 2016), and the adjusted p-value<0.01 & log2FoldChange>0 was considered significantly enriched.

For the datasets where raw data were unavailable, the reported HTSeq count table was used as input for DESeq2, and the conflicting transcript annotations between ENSEMBL releases were manually resolved. In the case of non-Illumina-based sequencing datasets, numerical log2foldchange was calculated from the reported TPMs of each gene. *R. norvegicus* datasets were converted to *M. musculus* genes using ENSEMBL BioMart annotations after the differential expression processing. All summary data obtained from our analysis is available in Supplementary Table 1.

Gene Ontology (GO) enrichment analyses were performed using a Bioconductor package TopGO (Alexa and Rahnenfuhrer, 2022). To prevent artefactual enrichment of brain-related terms, adult mouse brain transcriptome (Accession number: ENCFF468CXD) from the ENCODE project was used as background. Multiple testing was corrected via the Bonferroni method. An adjusted p-value<0.05 and foldchange>2 was considered a significant enrichment. Summaries of enriched GO terms were processed using the R package SimplifyEnrichment (Gu and Hübschmann, 2022) via clustering significantly enriched GO terms based on semantic similarity. Unclustered individual GO terms are given in Figure S2 and Supplementary Table 2

Conversion between human, mouse and fly orthologues (ENSEMBL release 99) was performed using the source data of DIOPT version 9 (Hu *et al*., 2011). To select for high-confidence orthologues, a cut-off score of 8 was used. Fly genes with the associated disease ontologies were acquired from FlyBase, and the enrichment of disease ontology terms were assessed with hypergeometric test followed by Bonferroni’s method multiple hypothesis correction.

To identify transcripts that are expressed in the three glial cell types (perineurial, subperineurial and ensheathing glial cells), single-cell nucleus sequencing data from Fly Cell Atlas was used (Li *et al*., 2022). The ‘Glial Cell’ loom file from 10x Cross-Tissue dataset was downloaded and filtered for non-glial cells based on the annotation. Using the R package ScopeLoomR (https://github.com/aertslab/SCopeLoomR), the digital gene expression matrices for the three cell types were extracted and the transcripts expressed in at least 5% of the corresponding clusters were identified.

For the Reactome pathway enrichment analysis, the gene to pathway query from Flymine was used to extract the Reactome ID annotations for each gene (https://www.flymine.org/flymine/templates/Gene_Pathway). The Reactome pathway hierarchical relationship between terms (available <u>here</u>) was used to group genes to the two highest hierarchical levels, the “top-level pathways” and the next hierarchical level, the “sub-pathways”, was used to calculate the enrichment.

All analysis scripts and data produced from this study are available at https://github.com/jefflee1103/Gala2023_glia-localised-RNA. The data that supports the findings of this study are available in the supplementary material of this article.

### 2.2 Experimental methods

#### 2.2.1 Fly stocks

The fly lines used in this research were raised at 25°C (all stocks used in the smFISH experiments and the spaced potassium assay experiment apart from *Nrg*-RNAi) or 30°C (larvae used for the *Nrg*-RNAi spaced potassium assay experiment in glial cells) on a standard cornmeal-agar food. To label glial cells in the smFISH experiments, a cross was made to obtain the following stock: UAS-mCD8-mCherry/CyoGFP; Repo>GAL4/Tm6B, Tb. This stock was then crossed to each of the CPTI lines and offspring selected for YFP and mCherry fluorescence. To label glia in the spaced potassium assay experiment with RNAi of transcripts of interest in glial cells, the following stock was constructed: UAS-mCD8-GFP/UAS-mCD8-GFP; Repo>GAL4/Tm6B, Tb. For each RNAi experiment, a homozygous line where a hairpin targets the coding sequence of the gene of interest with least number of off-targets was selected. If the progeny of the RNAi cross was embryonic or larval lethal, the next most suitable RNAi line meeting the above criteria was used. These lines were then crossed to the UAS-mCD8-GFP/UAS-mCD8-GFP; Repo-GAL4/Tm6B, Tb, and offspring selected for GFP fluorescence and lack of Tm6B, Tb phenotype. The control for the UAS-RNAi experiment was UAS-mCD8-GFP/+; Repo-GAL4/UAS-mCherry-RNAi. The glial cell specific knockdown of the 11 localized transcripts resulted in lethality for *Cip4*, where no 3rd instar Drosophila larvae of the correct genotype were observed. For Atpalpha (GD3093) and *Lac* (KK107450), the UAS-RNAi larvae were extremely few, small and sickly, and the experiments could not be performed on them. Using other UAS-RNAi lines for *Atpalpha* (HMS00703) and *Lac* (GD35524) allowed us to continue with experiments on those knockdown larvae. We observed no adults of the UAS-RNAi genotype for *nrv2*-RNAi, *Lac*-RNAi, *Cip4*-RNAi, and *Vha55*-RNAi. We observed adults for all other Repo>UAS-RNAi. crosses. **Table 1** lists all the strains used in the study.

**Table 1.** List of *Drosophila melanogaster* lines which were utilized in this project.

| Line name | Line ID | Line source | Line description |
| --- | --- | --- | --- |
| *nrv2*::YFP | CPTI001455 | Kyoto Stock Centre - *Drosophila* Genomics and Genetic Resources (DGRC) (https://www.dgrc.kit.ac.jp/) | *Nervana 2* gene YFP protein trap line ((Lowe *et al.*, 2014) |
| *Flo-2*::YFP | CPTI001427 | DGRC | *Flotillin 2* gene YFP protein trap line |
| *Cip4*::YFP | CPTI003231 | DGRC | *Cdc42-interacting protein 4* gene YFP protein trap line |
| *Vha55*::YFP | CPTI002645 | DGRC | *Vacuolar H<sup>+</sup>-ATPase 55kD subunit* gene YFP protein trap line |
| *Atpalpha*::YFP | CPTI002636 | DGRC | *Na pump $\alpha$ subunit* gene YFP protein trap line |
| *Nrg*::YFP | CPTI002761 | DGRC | *Neuroglian* gene YFP protein trap line |
| *Lac*::YFP | CPTI001714 | DGRC | *Lachesin* gene YFP protein trap line |
| *alpha-Cat*::YFP | CPTI002408 | DGRC | *$\alpha$ Catenin* gene YFP protein trap line |
| *Pdi*::YFP | CPTI002342 | DGRC | *Protein disulfide isomerase* gene YFP protein trap line |
| *gs2*::YFP | CPTI001918 | DGRC | *Glutamine synthetase 2* gene YFP protein trap line |
| Repo-GAL4 | RRID:BDSC_7415 | Bloomington Drosophila Stock Center (BDSC) (https://bdsc.indiana.edu/index.html) | Expresses GAL4 in glia |
| UAS-mCD8-mCherry/cyo-GFP | RRID:BDSC_27391 | BDSC | Expresses Cherry RFP fused to the mouse CD8 extracellular and transmembrane domains for membrane targeting under UAS control. |
| UAS-mCD8-GFP/cyo-GFP | RRID:BDSC_63045 | BDSC | Expresses GFP fused to the mouse CD8 extracellular and transmembrane domains for membrane targeting under UAS control. |
| cyo-GFP/Gla; tm6, tb/pri | Crossed from BDSC stocks | BDSC | A double balanced stock carrying Cyo-GFP over Gla on the 2<sup>nd</sup> chromosome and tm6, tb over prickles on the 3<sup>rd</sup> chromosome. |
| *Atpalpha*-RNAi | RRID:BDSC_32913 | BDSC | Expresses dsRNA for RNAi of *Atpalpha* under UAS control in the VALIUM20 vector. |
| *Atpalpha*-RNAi | GD3093 | Vienna Drosophila Resource Center (VDRC) | Expresses dsRNA for RNAi of *Atpalpha* under UAS control in the pUAST vector pMF3. |
| *alpha-Cat*-RNAi | KK107298 | VDRC | Expresses dsRNA for RNAi of *alpha-Cat* under UAS control in the pUAST vector pMF3. |
| *nrv2*-RNAi | GD960 | VDRC | Expresses dsRNA for RNAi of *nrv2* under UAS control in the pUAST vector pMF3. |
| *shot*-RNAi | RRID:BDSC_28336 | BDSC | Expresses dsRNA for RNAi of *shot* under UAS control in the VALIUM10 vector. |
| *Vha55*-RNAi | GD_9363 | VDRC | Expresses dsRNA for RNAi of *Vha55* under UAS control in the pUAST vector pMF3. |
| *Lac*-RNAi | GD_35524 | VDRC | Expresses dsRNA for RNAi of *Lac* under UAS control in the pUAST vector pMF3. |
| *Lac*-RNAi | KK_107450 | VDRC | Expresses dsRNA for RNAi of *Lac* under UAS control in the pUAST vector pMF3. |
| *Flo-2*-RNAi | VSH_330316 | VDRC | Expresses dsRNA for RNAi of *Flo-2* under UAS control in the WALIUM20 vector. |
| *gs2*-RNAi | GD_9378 | VDRC | Expresses dsRNA for RNAi of *gs2* under UAS control in the pUAST vector pMF3. |
| *Pdi*-RNAi | GD13418 | VDRC | Expresses dsRNA for RNAi of *Pdi* under UAS control in the pUAST vector pMF3. |
| *mCherry*-RNAi | RRID:BDSC_35785 | BDSC | Expresses dsRNA for RNAi of *mCherry* under UAS control in the VALIUM20 vector. |
| *nrv2*-GAL4 | RRID:BDSC_6800 | BDSC | Expresses GAL4 in wrapping glia. |
| 46F-GAL4 | - | Gift from Dr. Stefanie Schirmeier | Expresses GAL4 in perineurial glia (Xie and Auld, 2011). |
| *Mdr65*-GAL4 | RRID:BDSC_50472 | BDSC | Expresses GAL4 in subperineurial glia. |

#### 2.2.2 Solutions and reagents

The Haemolymph-Like Salines (HL3 solutions) were prepared as described previously (Roche *et al*., 2002; Ataman *et al*., 2008). The list of all the solutions used is given below (**Table 2**).

**Table 2.** A list of all the solutions used in the project.

| Solution | Composition |
| --- | --- |
| HL3 0.3 mM Ca<sup>2+</sup> | NaCl 70 mM, KCl 5 mM, MgCl<sub>2</sub> 20 mM, NaHCO<sub>3</sub> 10 mM, trehalose 5 mM, HEPES 5 mM, sucrose 115 mM, Ca<sup>2+</sup> 0.3 mM, pH 7.2 |
| HL3 1 mM $\text{Ca}^{2+}$ | NaCl 70 mM, KCl 5 mM, $\text{MgCl}_2$ 20 mM, $\text{NaHCO}_3$ 10 mM, trehalose 5 mM, HEPES 5 mM, sucrose 115 mM, $\text{Ca}^{2+}$ 1 mM, pH 7.2 |
| HL3 high $\text{K}^+$ | NaCl 40 mM, KCl 90 mM, $\text{Ca}^{2+}$ 1.5 mM, $\text{MgCl}_2$ 20 mM, $\text{NaHCO}_3$ 10 mM, trehalose 5 mM, HEPES 5 mM, sucrose 5 mM, pH 7.2 |
| 10x PBS | 1.37 M NaCl, 27mM KCl, 100mM $\text{Na}_2\text{HPO}_4$, 20mM $\text{KH}_2\text{PO}_4$, pH 7.4 |
| PBSTx | PBS 1x, 0.3% Triton X (v/v) |
| 20xSSC | 20g NaCl, 100.5g Tri-Sodium Citrate, pH 7.0 in 1L |
| TE | 10 mM Tris-HCL, 1mM EDTA, pH 8.0 |

#### 2.2.3 smFISH probes

Probes for the smFISH protocol were designed using a protocol described before (Gaspar, Wippich and Ephrussi, 2017). A set of oligonucleotides (28 for the YFP exon) against the gene region of interest was composed using LGC Biosearch Technologies’ Stellaris® RNA FISH Probe Designer. The oligonucleotides were pooled and elongated overnight at 37°C with a ddUTP conjugated to a desired dye (Atto 633 for the YFP probe) using terminal deoxynucleotidyl transferase enzyme from Life Technologies (Thermo Fisher Scientific). The fluorescently labelled oligonucleotides were then purified by Oligo Clean & Concentrator kit (Zymo Research®) and eluted in TE buffer, after which their concentration and degree of labelling were measured using a NanoDrop spectrophotometer. The probes were diluted with TE buffer to 25_μ_M concentration.

#### 2.2.4 RNA single molecule *in situ* hybridization (smFISH) on the *Drosophila* larval fillet

RNA single molecule in situ hybridization (smFISH) was carried out as described previously (Titlow *et al*., 2018). In short, wandering 3^rd^ instar (L3) larvae were dissected in HL3 0.3 mM Ca^2+^ as described before to produce a larval fillet with exposed NMJs (Brent, Werner and McCabe, 2009), fixed for 30 minutes at room temperature (RT) using 4% paraformaldehyde in PBS containing 0/1% Triton-X (PBSTx) and permeabilized 2x for 20 minutes in PBSTx at RT. Samples were then pre-hybridized for 20 minutes at 37°C in wash buffer (2x SSC, 10% formamide (Sigma-Aldrich, F9037)), and then hybridized overnight at 37°C in hybridization buffer (10% formamide, 10% dextran sulphate (J62787.18, Alfa Aesar), 250nM smFISH probe(s), anti-HRP (**Table 3**) in 2x SSC). Samples were then rinsed in wash buffer again and counterstained with DAPI (1:1000 from 0.5mg/mL stock) in wash buffer for 45 minutes at RT. Next, samples were washed for 45 minutes in wash buffer at RT and incubated in Vectashield anti-fade mounting medium (Vector Laboratories) adjusted for the objective refractive index for 30 minutes, and subsequently mounted on the slide in the mounting medium.

**Table 3.** A list of all the primary antibodies used in this study.

| Antibody | Detects | Source | Species | Dilution |
| --- | --- | --- | --- | --- |
| Anti-Dlg1 | Disks large, a post-synaptic scaffolding protein | Developmental Studies Hybridoma Bank, https://dshb.biology.uiowa.edu/ | Mouse | 1:200 |
| Anti-HRP conjugated to Alexa 405, 488, 568 or 647 | Neurons | Jackson ImmunoResearch Europe Ltd | Goat | 1:100 |

#### 2.2.5 Immunofluorescence (IF) on the *Drosophila* larval fillet

L3 larvae were dissected and fixed as described in the smFISH protocol. Larvae were then blocked for more than 1 hour at 4°C in blocking buffer (PBSTx, 1.0% BSA). Samples were incubated overnight at 4°C with the primary antibody (**Table 3**) in blocking buffer. The next day, samples were washed for about 1 hour and incubated in the secondary antibody solution (conjugated to Alexa Fluor 488, 568 or 647, used at 1:500, from Life Technologies, diluted in PBSTx) together with DAPI (1:1000 from 0.5mg/mL stock) for a further 1 hour. Samples were then washed for 45 minutes in PBSTx at RT and incubated in Vectashield and mounted as in the smFISH protocol (section 2.2.4).

#### 2.2.6 Spaced High K^+^ depolarization paradigm

The spaced potassium assay has been carried out as described before (Piccioli and Littleton, 2014; Roche *et al*., 2002; Ataman *et al*., 2008). Briefly, the larvae were dissected in 0.3 mM Ca^2+^ HL3 and then moved to 1 mM Ca^2+^ HL3. The unstretched larvae were then washed with high K^+^ HL3 for the periods of 2, 2, 2, 4 and 6 minutes with 15 minutes intervals of 1 mM Ca^2+^ HL3 in between (for RNAi knockdown experiments on *Atpalpha*, *alpha-Cat*, *nrv2*, *shot*, *Nrg*, *Flo-2*, *Vha55*), or periods of 2 minutes, three times, with 10 minutes in between (for RNAi knockdown experiments on *Lac*, *gs2*, *Pdi*). The assay was performed with a minimum of 5 control and 5 RNAi larvae per each experiment and in one replicate for knockdown of *Atpalpha*, *alpha-Cat*, *Vha55*, two replicates for knockdown of *Nrg* and *Flo-2*, six replicates for the knockdown of *Lac*, three replicates for all remaining RNAi knockdown experiments. In each experiment, an internal control was performed at the same time, where the Repo>mCD8-GFP larvae were crossed to UAS-RNAi line against mCherry protein, absent in these larvae, and each experiment was compared to its internal control only. After the spaced potassium pulses, the larvae were then stretched again, left for a period of rest of 30 minutes, and fixed and labelled as described in the IF protocol. For all experiments, segments A2-A5 of muscles 6/7 were imaged.

#### 2.2.7 Image acquisition and processing

For smFISH experiments, the larvae were dissected, fixed and stained with DAPI and anti-HRP antibody conjugated to the DyLight 405 dye, and the anti-YFP exon probe conjugated to Atto-633 dye, as per the smFISH protocol. A minimum of 3 larvae and 15 NMJs were assessed. For the potassium stimulation experiments, a minimum of 5 control and 5 RNAi larvae were dissected, treated as described above (see “Spaced High K^+^ depolarization paradigm”) fixed and stained with DAPI and anti-HRP antibody conjugated to Alexa-647 dye, and the anti-Dlg1 antibody as described in the IF protocol, and the glial membrane was labelled in endogenously with Repo>mCD8-GFP. Mounted specimens were imaged using an inverted Olympus FV3000 laser scanning confocal microscope or Olympus CSU-W1 SoRa laser spinning disk confocal microscope. Images were acquired using 60x 1.4NA Oil UPlanSApo objective (FV3000) or 100x 1.51NA Oil UPlanSApo objective, 100x 1.4NA Oil UPlanSApo objective or 60x 1.51NA Oil UPlanSApo objective (SoRa). Laser units were solid state 405, 488, 568 and 640 lasers for both microscopes.

The images were then processed in ImageJ (https://imagej.nih.gov/ij/) (Schneider, Rasband and Eliceiri, 2012). The glial processes at the NMJ are largely flat, hence the 2 dimensional areas of the GFP labelled glial membranes were measured as previously described; these membranes included small sections of non-synaptic motor axon branches from full Z-stack 2D projections as well as glial membrane nearing the synaptic lamella (Brink *et al*., 2012b). The neurite areas were measured identically, except they were labelled with anti-HRP. The settings used were: the linear auto-contrast, auto-threshold, and area measurement functions of NIH ImageJ (Abràmoff, Magalhães and Ram, 2004). In the spaced potassium stimulation experiment, the ghost boutons were quantified manually.

For statistical analysis, R Studio was used. Each two-dimensional area from each NMJ at muscles constituted an independent replicate (“n”) value. The n numbers for each genotype and experiment are specified in respective figure legends. For spaced potassium stimulation experiment, we quantified and reported average log2 foldchange of bouton counts post potassium stimulation compared to RNAi controls and performed the Wilcoxon rank sum test. For the areas’ quantification, we calculated foldchange in glial protrusion area, neurite area and their ratio upon knock-down of glial protrusion-localized transcripts. We performed these calculations in two sets: before and after potassium activation assay. For each set, average foldchange for each gene was calculated. Student’s t-tests were performed on the data to assess significant differences. The summary statistics are available in Supplementary Table 1 (unstimulated NMJs areas), Supplementary Table 1 (stimulated NMJs areas) and Supplementary Table 3 (stimulated NMJs ghost boutons).

## 3 RESULTS

### 3.1 Collation of multiple studies characterizing transcript localization to peripheral glial cytoplasmic projections

To identify a common set of localized glial transcripts, we performed a meta-analysis of, to our knowledge, all published mammalian localized and synaptic transcriptomic datasets derived from multiple types of glia (Thomsen *et al*., 2013; Thakurela *et al*., 2016; Boulay *et al*., 2017; Sakers *et al*., 2017; Azevedo *et al*., 2018; Mazaré *et al*., 2020; Vasek *et al*., 2021), outlined in **Table 4**. Our first goal was to integrate all the raw data, make it interoperable and collate it into a unified data set that can be interrogated as a single entity.

**Table 4.**
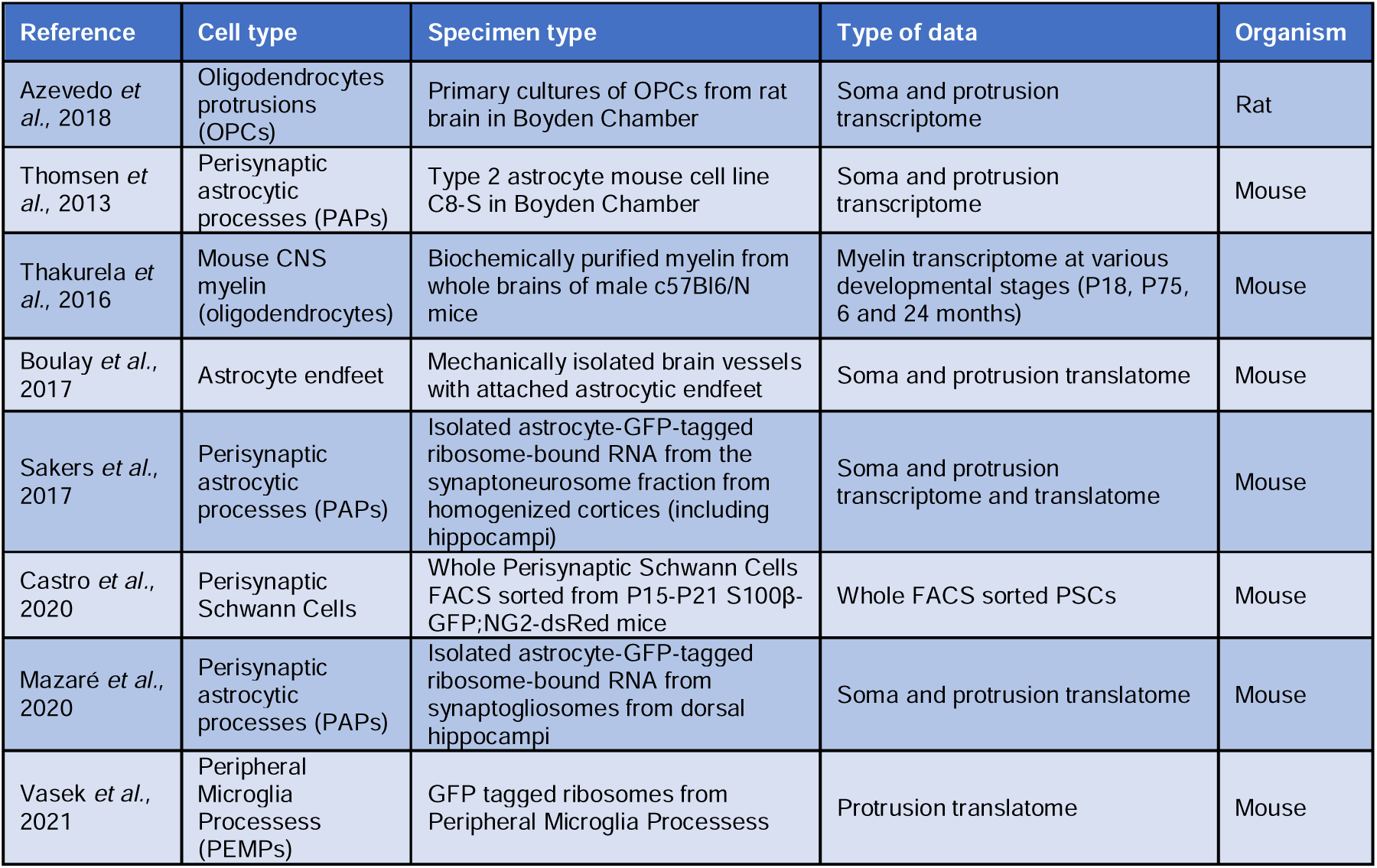
A summary of all the available localized and synaptic glial transcriptomic datasets which were analyzed in this study. Most of the datasets explored transcripts localized to glial periphery, but the PSCs dataset was also included although it involved bulk transcriptomics because of its relevance to the glial involvement in synaptic plasticity.

These glial cell types include four studies of astrocytes, and one study each of oligodendrocytes, micro-dissected myelin and microglia. We also included one bulk transcriptomic study from Perisynaptic Schwann cells (PSCs) that are non-myelinating glia associated with the NMJ that were isolated from mice. We used the PSCs study because the isolated transcripts are associated with synapse activity, and therefore the dataset is highly relevant in our quest for mechanistic connections between mRNA localization and synaptic plasticity (Castro *et al*., 2020). The 8 studies used various methods of purifying localized transcripts, which are summarized in Table 4 “Specimen type” column, and in Figure 1A. We collated 8 transcriptomic and 4 TRAP (Translating Ribosome Affinity Purification) libraries, using a standardized pipeline for interoperability between the 12 datasets (Figure 1B). For details of how our analysis was performed and how we created a single data framework from all these independently acquired data sets, see the “Data meta-analysis methods” section of the “Materials and Methods”.

**Figure 1.**
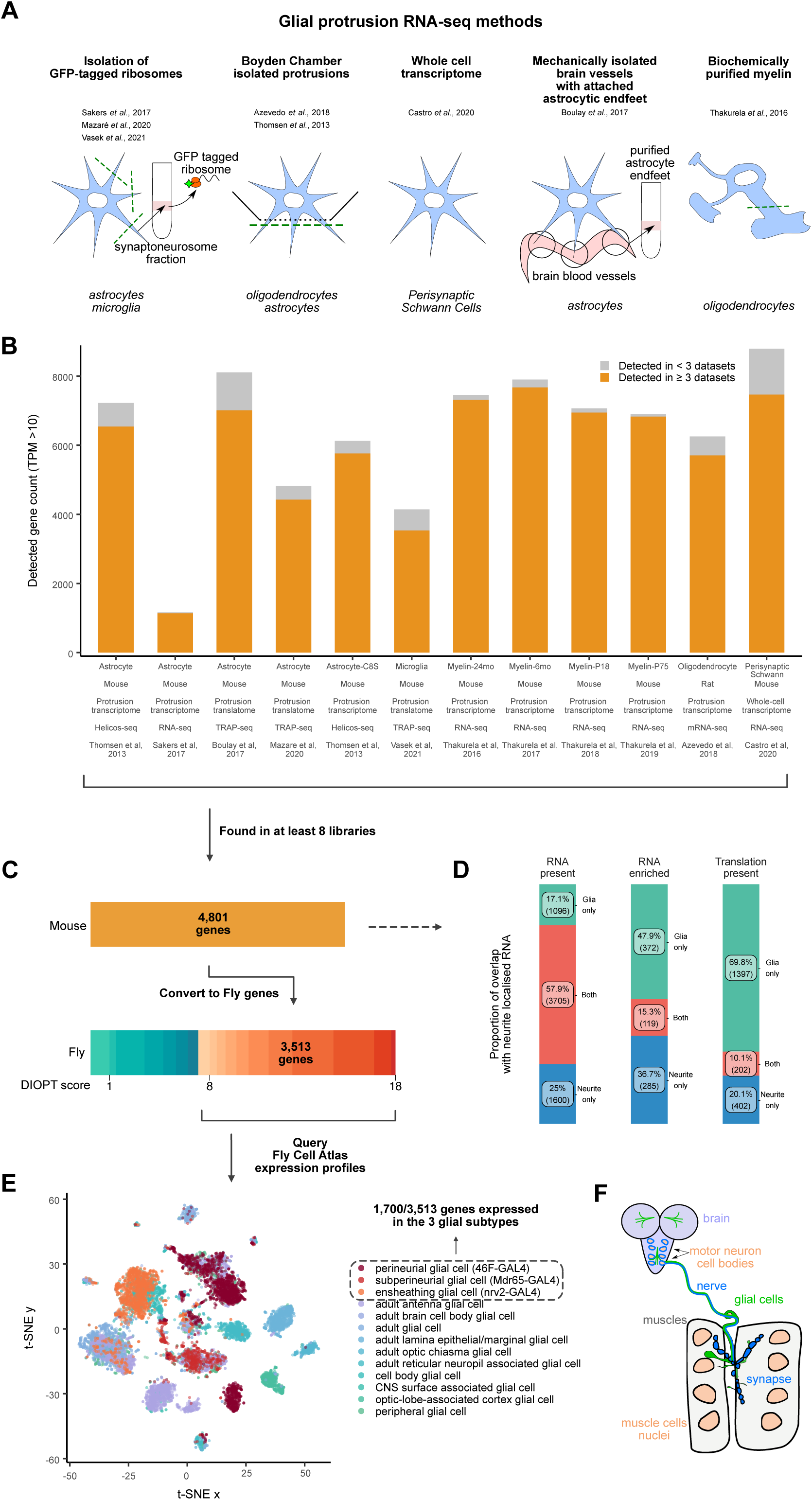
Conserved transcripts localized to glial protrusions. A) A graphical representation of the techniques used to separate the protrusion-localized transcripts in each study, with the exception of PSCs (middle), where whole cells were used. B) Bar graph showing the datasets included in this study. High confidence expression cut-off was set at TPM > 10, and transcripts detected in more than 3 independent datasets are shown in orange color. C) Identification of high confidence *D. melanogaster* orthologs of 4,801 genes that were detected in at least 8 datasets. DRSC Integrative Ortholog Prediction Tool (DIOPT) score of 8 was used as cut-off. D) Comparison of transcripts localized, enriched or translated in the glial protrusion and/or neurites (von Kügelgen and Chekulaeva, 2020). Transcripts were considered enriched when log2FoldChange > 0 and adjusted p-value < 0.05, comparing protrusion and soma compartment-specific RNA-seq libraries. TRAP-seq TPM > 10 was considered as existence of translation. E) t-SNE plot of combined glial single-nuclei RNA-seq from the Fly Cell Atlas (Li *et al*., 2022). Perineurial, subperineurial and ensheathing glial cell clusters are depicted in orange-red colors. F) A schematic representing the distribution and location of glial cells in the *Drosophila* 3^rd^ instar larva. Six subtypes of glia exist in the larvae: perineurial and subperineurial glia, cortex glia, astrocyte-like and ensheathing glial cells and, finally, the peripheral nervous system (PNS) specific wrapping glia (Yildirim *et al*., 2019). Ensheathing glia of the CNS are continuous with the wrapping glia of the CNS. The perineurial and subperineurial glia have long and extensive projections which reach all the way to the neuromuscular junction (NMJ).

Using our bespoke data processing pipeline, we found 4,801 common transcripts detected in at least 8 libraries of localized mammalian glial transcripts. We tested whether this set of transcripts represents a population of localized mRNAs unique to the periphery of glia or whether they are also shared with the neurite transcriptome. To distinguish these possibilities, we compared the combined list of glial localized transcripts to the group of transcripts shown to commonly localize to the mammalian neuronal projections (von Kügelgen and Chekulaeva, 2020). We found a large overlap between the transcripts present at the periphery of both glia and neurons (Figure 1D). Our observations agree with a recent study highlighting conserved mRNA transport mechanisms between cell types (Goering, Arora and Taliaferro, 2022) and significant overlap of localized transcripts within astrocytes, radial glia and neurons (D’Arcy and Silver, 2020).

### 3.2 Meta-analysis of localized glial transcriptomes yields a predicted group of transcripts localized to the periphery of *Drosophila* peripheral glia

We then asked whether the mammalian glial localized transcripts were likely to be conserved across higher eukaryotes. To address this question in a specific case that is highly experimentally tractable, we intersected our mammalian dataset with the glial subtypes that extend their processes to the Peripheral Nervous System (PNS) of *Drosophila*, a well-established model for studying mRNA transport and local translation. We first filtered the 4,801 mammalian genes to only include the ones that have high confidence *Drosophila* homologues, retaining genes with a DIOPT score of at least 8 as a cutoff (Figure 1C). This filtering process resulted in a list of 3,513 *Drosophila* genes that we used to query the glial nuclear transcriptome from the Fly Cell Atlas (Li *et al*., 2022) (Figure 1E). We chose transcripts that were expressed in perineurial glia (PG) and subperineurial glia (SPG), the glial subtypes known to extend their processes to the *Drosophila* Neuromuscular Junction (NMJ) (Figure S1B and S1C). These glia have very long cytoplasmic extensions, and their cell bodies can be hundreds of micrometers away from their furthermost projections (Brink *et al*., 2012a; Sepp, Schulte and Auld, 2000; Sepp, Schulte and Auld, 2001; Sepp and Auld, 2003) (Figure 1E). We also included ensheathing glia (EG), as they are the closest glial cell type in the Fly Cell Atlas that corresponds to wrapping glia (WG) of the PNS. The ensheathing glia in the CNS are continuous with the wrapping glia in the PNS, and express similar marker genes such as *nrv2* (Yildirim *et al*., 2019) (Figure S1A). Our filtering of the mammalian glial localized mRNAs with high confidence *Drosophila* homologues by expression in any of the three *Drosophila* PNS glial subtypes (PG, SPG and WG/EG) yielded a group of 1,700 transcripts that we classify as “predicted to be present” in the projections of the *Drosophila* PNS glia. We hereafter refer to these transcripts as localized, in which we put emphasis on the presence of transcripts in the distal periphery rather than their enrichment compared with the cell body.

### 3.3 Gene ontology analysis reveals the statistical enrichment of transcripts related to mRNA trafficking, membrane composition and cytoskeletal regulation

To understand the functional characteristics of this group of localized transcripts, we identified enriched Gene Ontology (GO) terms followed by semantic similarity clustering of related terms within the localized 1,700 genes (Figure 2). Unclustered individual GO term enrichment results are given in Figure S2 and Supplementary Table 2. Our analysis revealed that the localized transcripts are enriched for biological processes involving morphogenesis/development, subcellular localization, cytoskeletal organization, signaling processes, and mRNA metabolism (Figure 2A). In terms of molecular function, RNA-binding, cytoskeleton-binding, as well as transmembrane transporter genes were highly enriched in our gene set (Figure 2B). Finally, our analysis of GO terms for the cellular component showed enrichment of terms related to vesicle/membrane transport, ribonucleoprotein granules and ribosomes in connection with the cytoskeleton, and the synapse, including synaptic terminal, boutons and junctions (Figure 2C). Taken together, our GO enrichment analysis highlights statistically significant over-representation of functions related to membrane trafficking, cytoskeleton regulation, local translation, and cell-cell communication within the localized transcripts, all of which are likely to be very active at the distal periphery of polarized cells.

**Figure 2.**
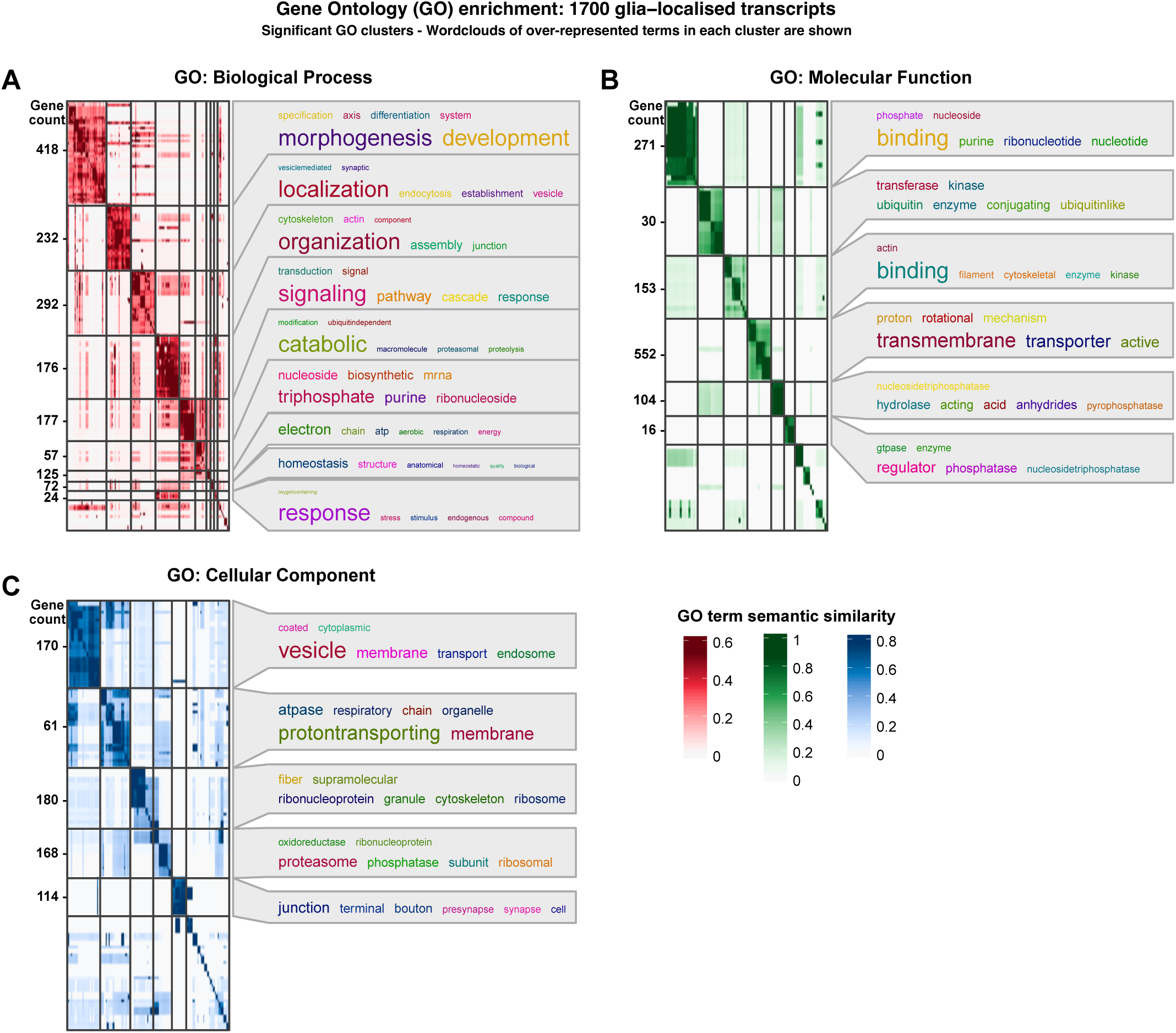
Functional annotation of glial protrusion-localized transcripts. A-C) Overviews of gene ontology (GO) terms enriched in the 1,700 glial protrusion-localized transcripts Biological Process, Molecular Function and Cellular Component categories. Enriched GO terms (FoldChange > 1.5, adjusted p-value < 0.01) were clustered based on the semantic similarity. Word clouds for each cluster show over-represented terms. *D. melanogaster* genes with high-confidence orthologues to *M. musculus* genes (DIOPT score ≥ 8) were used as background.

### 3.4 The 1,700 transcripts predicted to be present at the glial periphery are statistically enriched in neurodegenerative and neuropsychiatric disorders

Neurodegenerative and neuropsychiatric disorders have been associated with disruption of a number of post-transcriptional mechanisms, such as local translation, RNA-binding protein (RBP) activities and formation of RNA-rich granules (Blanco-Urrejola *et al*., 2021). To determine whether our predicted *Drosophila* glial peripherally localized transcripts are statistically enriched in associations with previously studied nervous system disorders, we carried out a disease ontology enrichment analysis. We found a significant enrichment of disease terms related to neuropathologies in our set of 1,700 glial protrusion-localized transcripts (Figure 3A). Interestingly, terms related to neurodegeneration and dementia were particularly enriched in the 1,700 genes and were also found in previous studies to be connected to glial related mechanisms that cause the diseases (Blanco-Urrejola *et al*., 2021). We also compared the set of 1,700 glial protrusion-localized transcripts with the SFARI gene database, which is a well-annotated list of genes that have associations with autism spectrum disorder (https://www.sfari.org/). We found a significant overlap between the 1,700 glial protrusion-localized transcripts and the SFARI genes (Figure 3B), further supporting the idea that mRNA localization in glia and the specific list of 1,700 genes we have highlighted are statistically enriched in genes associated with neurological disorders.

**Figure 3.**
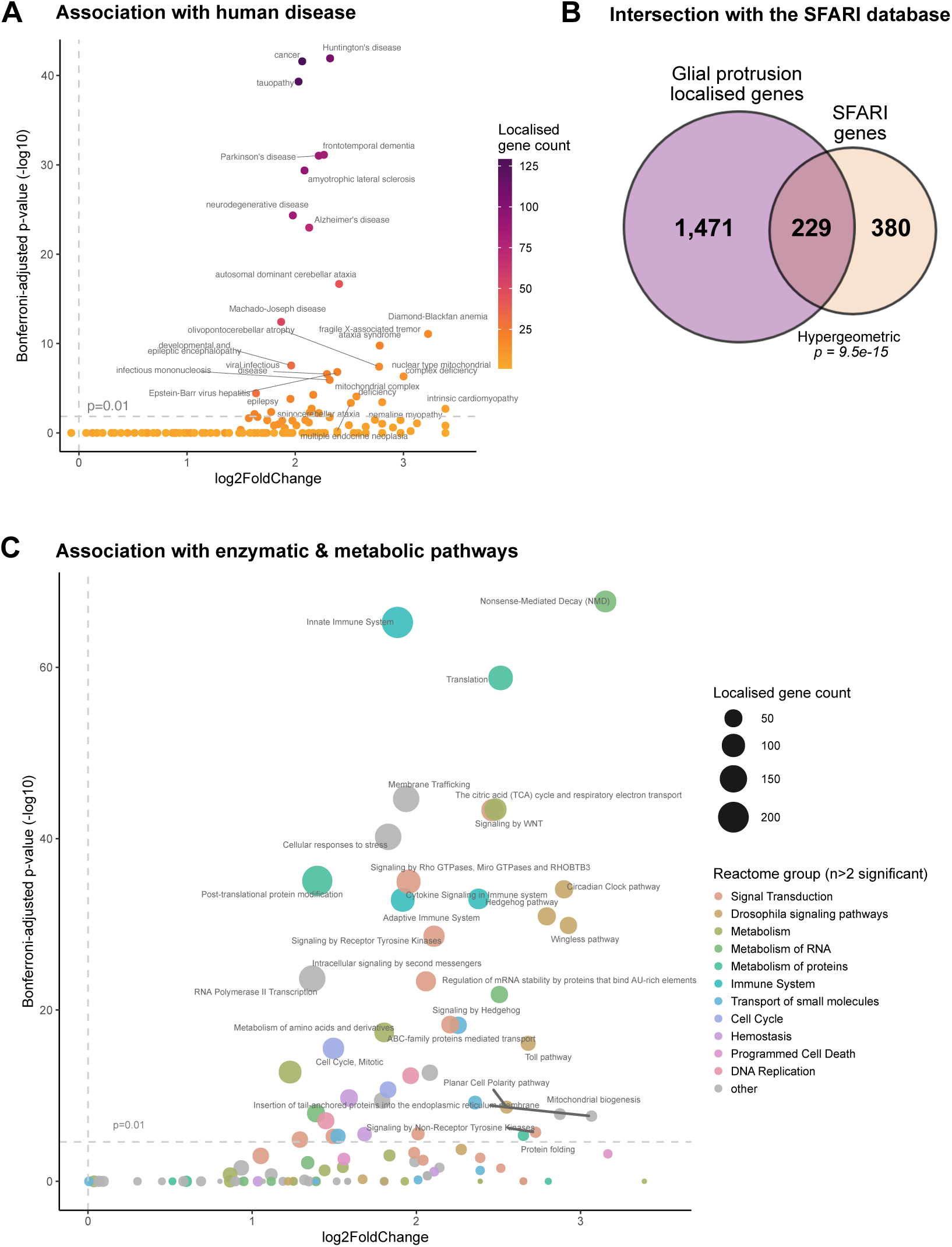
Association of glial protrusion-localized transcripts with disease. A) Enrichment of disease ontology (DO) terms in the 1,700 glial protrusion-localized transcripts. DO terms from *D. melanogaster* genes with high-confidence orthologues to *M. musculus* genes (DIOPT score ≥ 8) were used as background. Enrichment was assessed with hypergeometric test and corrected for multiple hypothesis testing. **B)** Significant overlap between the 1,700 glial protrusion-localized transcripts and the SFARI list of genes. SFARI gene is an annotated list of genes that have been investigated in the context of autism spectrum disorder (Banerjee-Basu and Packer, 2010). 1,095 *H. sapiens* SFARI genes were converted to *D. melanogaster* genes (DIOPT score ≥ 8) before the statistical enrichment. **C)** Reactome pathway enrichment analysis of the 1,700 glial protrusion-localized transcripts against the whole D. Melanogaster genome as background. The size of the of the points shows the number of localized genes corresponding to the term and the color of the point indicates the parent term in the Reactome pathways hierarchy. Enrichment was assessed with hypergeometric test and corrected for multiple hypothesis testing.

It was also very interesting to determine whether the 1,700 localized glial transcripts we highlighted are also enriched in signaling and enzymatic pathways. To investigate this possibility, we carried out a reactome-pathway enrichment analysis and found a significant enrichment of terms related to signaling pathways, such as Hedgehog or Wnt pathways, as well as terms related to mRNA translation, mRNA stability regulation and nonsense-mediated decay (Figure 3C). These significant associations led us to hypothesize that many of the localized mRNAs at the periphery of glia are required for neuron-glia, glia-muscle or glia-glia communication, and localized mRNA processing and metabolism at the distal cytoplasm of glia. Such mechanisms could potentially include non-canonical mRNA processing that has been previously suggested to be involved in brain physiology and cancer (Pitolli *et al*., 2022).

### 3.5 More than 70% of the previously identified glial transcripts are predicted to be present in the *Drosophila* PNS glia

Do the predicted transcripts indeed localize to the periphery of glial cells? To test our prediction in *Drosophila* glia, we cross-referenced the 1,700 transcripts against a list of 19 transcripts, which we previously identified experimentally to be present in the *Drosophila* peripheral glia by screening 200 randomly chosen genes for their localization status throughout the nervous system, including NMJ glia (Titlow *et al*., 2022). We found that 15 out of 19 fly genes had high confidence mammalian homologues, so we carried out our comparison only with these 15 genes (see Table 5).

**Table 5.** A table indicating whether the 19 genes previously predicted to be present in the Drosophila glia have been predicted in the list of 1,700 Drosophila homologs. All transcripts which did not have a high confidence D. melanogaster orthologs of 4,801 genes that were detected in at least 8 datasets (DIOPT score ≥ 8) were removed.

| Found in Titlow et al. | FBgn ID | Is predicted in 1,700? |
| --- | --- | --- |
| *Lac* | FBgn0010238 | Yes |
| *Pdi* | FBgn0286818 | Yes |
| *nrv2* | FBgn0015777 | Yes |
| *Lost* | FBgn0263594 | No |
| *Flo-2* | FBgn0264078 | Yes |
| *alpha-Cat* | FBgn0010215 | Yes |
| *Vha55* | FBgn0005671 | Yes |
| *Atpalpha* | FBgn0002921 | Yes |
| *Nrg* | FBgn0264975 | Yes |
| *Cip4* | FBgn0035533 | Yes |
| *Nrx-IV* | FBgn0013997 | No |
| *gs2* | FBgn0001145 | Yes |
| *shot* | FBgn0013733 | Yes |
| *kst* | FBgn0004167 | No |
| *sdk* | FBgn0021764 | No |

We found that 11 out of the 15 transcripts (73.33%) were predicted from our analysis of mammalian glial transcripts to localize to the glial periphery in *Drosophila*. As these data originate from a systematic study of 200 mRNAs across the larval nervous system (Titlow *et al*., 2022), we sought to repeat the experiments testing the localization status of the 11 genes in greater detail, using smFISH together with markers of specific glial sub-types (Figure 4 and S4, see “Experimental methods” for details). Our results confirmed with greater confidence that all 11 transcripts were indeed present in the peripheral glia at the *Drosophila* NMJ. To determine the statistical significance of the 11/15 hits, we carried out an enrichment test that follows a hypergeometric distribution (independent sampling). Our results showed a slim probability of 8.37×10^-05^ for the 73% overlap to have occurred by chance, hence assuming a significant over-representation. We conclude that the predicted group of 1,700 are highly likely to be peripherally localised in *Drosophila* cytoplasmic glial projections.

**Figure 4.**
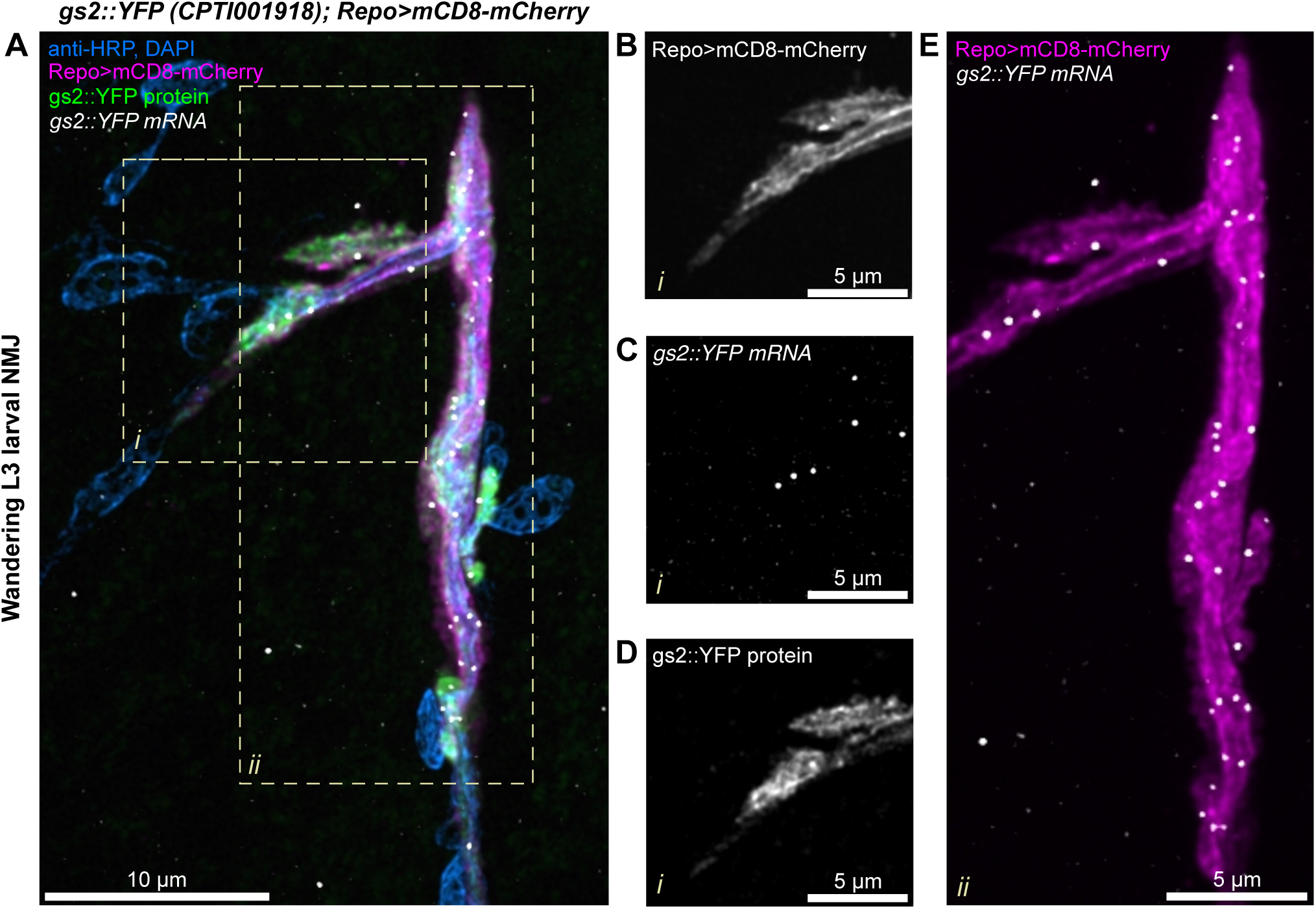
Localization of *gs2*::YFP transcripts in *Drosophila* glia. A) A confocal image of the *Drosophila* 3^rd^ instar larva NMJ (Segment A4), showing the neuron and muscle cell nuclei in blue (anti-HRP antibody conjugated to Alexa-405 fluor and DAPI, respectively), *gs2*::YFP protein in green, glial membrane of perisynaptic glia labelled with Repo>mCD8-mCherry in magenta, and the YFP exon associated with *gs2*::YFP protein in white (Atto-633). **B-D)** A single-channel view (see label) of a zoomed-in area **i)** of A) in which multiple mRNA molecules are present in the glial projection. **E)** A zoomed in area **ii)** of A) showing the entire length of the perisynaptic glia at that synapse with the mRNA molecules distributed regularly throughout the glial cell projections.

Our secondary analysis also allowed us to obtain higher resolution single mRNA molecule information on the distributions of mRNAs molecules across the motoneuron axons and the surrounding glial protrusions (Figure 4 and Figure S4). We found specific examples of transcripts that were highly enriched in glia, with a much lower abundance in the motoneurons, including *gs2*, *nrv2*, *flo-2* and *Lac* (Figure 4 and S4A, B, G respectively). Other transcripts were uniformly distributed in motoneurons, glia and muscle cells in the NMJ, including *Cip4*, *Pdi*, *Vha55*, *Nrg*, *alpha-Cat*, *Atpalpha*, *shot* (Figure S4, C-F, H-J, respectively). In conclusion, these results show that the localization of transcripts, extrapolated from vertebrates to *Drosophila,* were highly concordant, reliably predicting the presence of multiple mRNAs at the glial periphery and highlighting a possible evolutionary conservation of glial protrusion-localized transcriptome.

### 3.6 Localized glial transcripts influence plasticity in the adjacent motoneurons

To test the function of the glial localized mRNAs to influence synaptic plasticity of the neighboring motoneurons, we knocked down the genes specifically in glia, with the motoneurons remaining unaffected. We used 13 UAS-RNAi lines (against 11 different transcripts) driven by the pan-glial Repo-GAL4 driver (see materials and methods for further details). We assayed the impact of knocking down each gene in glia on the ability of the wild type NMJ motoneurons to make new synapses in response to chemical stimulation of the nerve. We applied the well-established protocol for activity-dependent synaptic plasticity through spaced pulses of potassium applied to a living NMJ preparation (Ataman *et al*., 2008; Roche *et al*., 2002; Piccioli and Littleton, 2014; Vasin *et al*., 2014). We found that 10 of the UAS-RNAi lines were viable until 3^rd^ instar larvae without causing excessive lethality.

RNAi knockdown of *Cip4* in glia did not yield viable 3^rd^ instar larvae, so this gene was excluded from further analysis. Therefore, we tested 10 out of 11 localized transcripts in glia for their involvement in regulating synaptic plasticity of motoneurons. Interestingly, we found that Repo>*Vha55*-RNAi larvae display a small central brain phenotype (Figure S5), however the larvae were otherwise indistinguishable in size from their control counterparts, so they were assayed as with other genotypes.

We performed the spaced potassium stimulation assay (Figure 5A) to quantitate the effect of glial-specific RNAi by counting the number of newly formed neuronal boutons defined as “ghost” boutons, which are immature structures that lack post-synaptic density markers such as Discs large 1 (*Dlg1*) protein (Ataman *et al*., 2008; Roche *et al*., 2002). We found that for 3 of the 10 transcripts (*Lac*, *gs2*, *Pdi*) the RNAi knockdown in *Drosophila* glia causes a reduction in the number of ghost boutons formed in response to pulses of potassium (Figure 5B, C). We conclude that *Lac*, *gs2* and *Pdi* are required within glia for the correct plasticity of the motoneuron synapses.

**Figure 5.**
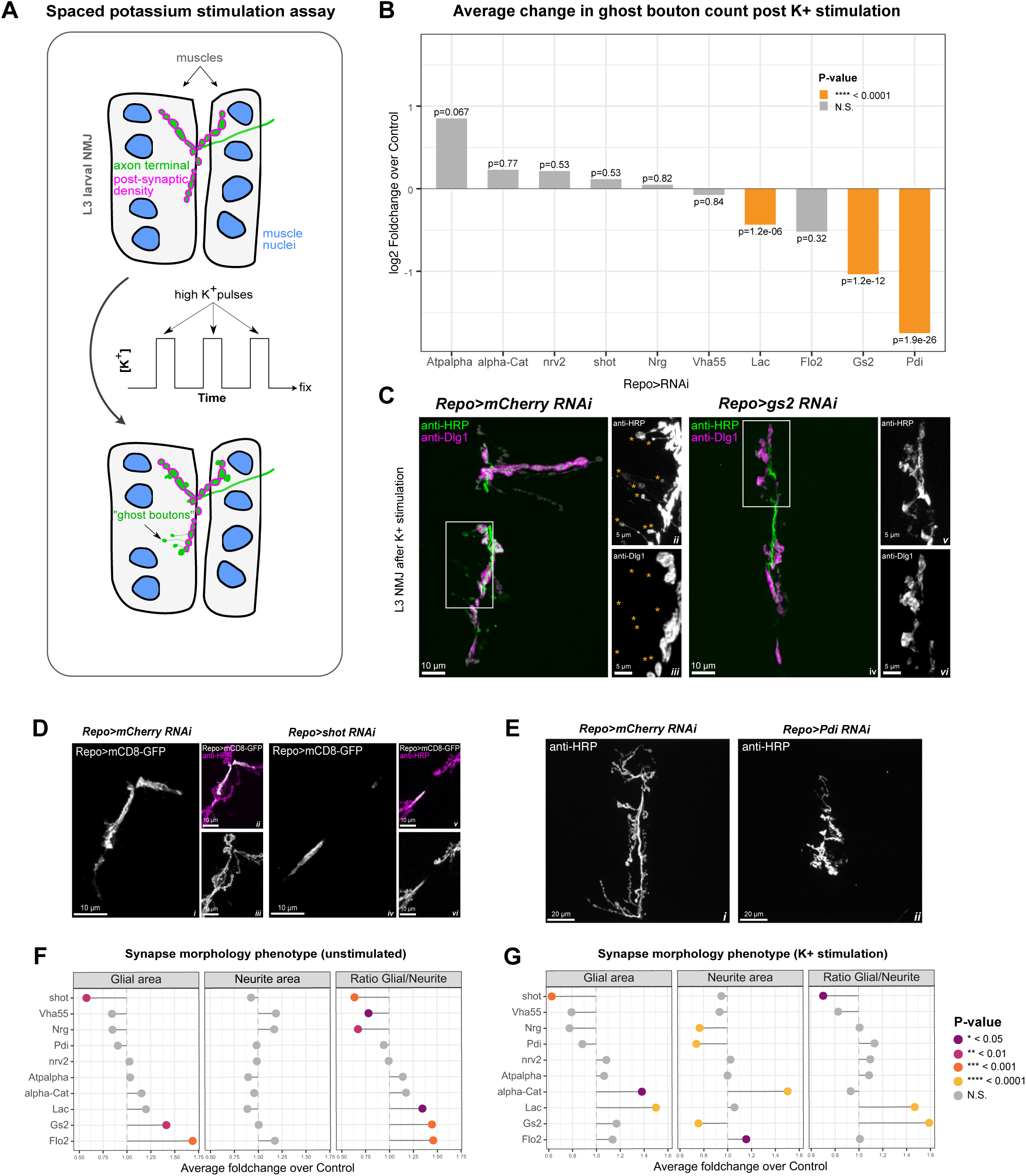
Knockdown of glial localized transcripts interferes with synaptic plasticity and development. A) A schematic representation of the spaced potassium stimulation assay used in this study. Larvae are dissected and subjected to pulses of high potassium solution mimicking neuromuscular stimulation. After the assay has been completed, newly formed axon terminal endings with immature synapses, called “ghost boutons”, can be detected using anti-HRP antibody labelling thanks to the lack of the post-synaptic density present, which can be labelled using anti-Discs-large antibody, which the ghost boutons lack. **B)** Knock-downs of multiple glial protrusion-localized transcripts disrupt synaptic plasticity. Bar graph represents average log2 FoldChange of bouton counts post potassium stimulation compared to RNAi Controls. Statistically significant changes are highlighted in orange color. Wilcoxon rank sum test p-values are reported. N = 30(*alpha-Cat*-RNAi), 30(*Atpalpha*-RNAi), 58(*Flo-2*-RNAi), 28(*Vha55*-RNAi), 98(*gs2*-RNAi), 238(*Lac*-RNAi), 48(*Nrg*-RNAi), 73(*nrv2*-RNAi), 114(*Pdi*-RNAi), 76(*shot*-RNAi) NMJs) **C)** Confocal images showing representative control **(i-iii)** and *gs2*-RNAi **(iv-vi)** synapses from the spaced potassium assay experiment. Axon terminal and ghost boutons are labelled in green, and the anti-Discs-large antibody labelling in magenta in i) and iv). ii) and iii) represent the area inside the white rectangle in i) (see in-image labels for details). v) and vi) represent the area inside the white rectangle in iv). ii) and iii) show multiple ghost boutons lacking the post-synaptic density for the Control panel indicated with orange asterisks. v) and vi) show an extreme case where no ghost boutons were found for that NMJ is presented for the *gs2*-RNAi panel. **D)** Confocal images showing representative control (i-iii) and *shot*-RNAi (iv-vi) synapses (see in-image labels for details). NMJ glial projections for *shot*-RNAi (iv) are much less expansive than the control in the unstimulated NMJs (i). **E)** Confocal images showing representative control (i) and *Pdi*-RNAi (ii) synapses (see in-image labels for details). The NMJ areas are much smaller for the *Pdi*-RNAi NMJs when compared to the control in the stimulated NMJs. **F-G)** Foldchange in glial protrusion area, neurite area and their ratio upon knock-down of glial protrusion-localized transcripts before (F, N = 28(*alpha-Cat*-RNAi), 30(*Atpalpha*-RNAi), 30(*Flo-2*-RNAi), 23(*Vha55*-RNAi), 30(*gs2*-RNAi), 28(*Lac*-RNAi), 23(*Nrg*-RNAi), 28(*nrv2*-RNAi), 28(*Pdi*-RNAi), 27(*shot*-RNAi) NMJs)) and after (G, N = 30(*alpha-Cat*-RNAi), 30(*Atpalpha*-RNAi), 58(*Flo-2*-RNAi), 28(*Vha55*-RNAi), 97(*gs2*-RNAi), 239(*Lac*-RNAi), 48(*Nrg*-RNAi), 73(*nrv2*-RNAi), 114(*Pdi*-RNAi), 79(*shot*-RNAi) NMJs)) potassium activation assay. Data represents average foldchange for each gene. Student’s t-test.

We also assayed the effect of knocking down the localized transcripts in glia on the morphology of the glial projections and of the wild-type axon terminals in the NMJ. We examined the morphologies of 3^rd^ instar glia expressing RNAi and wild type motoneurons (Figure 5 D, E) with or without potassium stimulation (Figure 5 F, G). We found that, even in the absence of stimulation, the knock-down of many of the genes resulted in aberrant growth or shrinkage of glial projections as well as a change in the ratio of glial to neuron surface areas, suggesting a developmental defect. For example, RNAi against *shot* resulted in near elimination of the glial projections. While in the control larvae, the glial projections extend on the surface of the muscle cells, in *shot*-RNAi larvae the non-synaptic motor axon branches are covered with glia, but the glia lack any projections onto the surface of the muscle (Figure 5D, E). We also saw varying defects in the sizes of both glial and neuronal projections after spaced potassium stimulation (Figure 5F, G). Our results demonstrate that many of the localized glial transcripts are required for the correct morphology of the *Drosophila* NMJ, and 30% are specifically necessary in glia for the correct synaptic plasticity of their neighboring neuron.

## 4 Discussion

### 4.1 Transcripts localized to the projections of different glial subtypes have conserved roles in coordinating synaptic plasticity and development

Using meta-analysis of multiple published datasets of peripherally localized mammalian glial mRNAs, we have predicted 1,700 localized transcripts in 3 glial subtypes associated with the *Drosophila* motoneurons and NMJ synapses. These mRNAs are highly enriched in *Drosophila* homologues of human genes with associations in a variety of neurodegenerative and neuropsychiatric diseases, which also often coincide with molecular and cellular functions that have known associations with the cell periphery (Figure 1). The localized glial mRNA we predict include regulators of cytoskeleton dynamics and remodeling, membrane dynamics, signaling pathways and mRNA metabolism (Figure 2). We tested some of these predictions by using smFISH to characterize in more detail the localization of 15 out of 19 of the mRNAs that we previously identified as present in *Drosophila* glia (Titlow *et al*., 2022) and possessing high confidence mammalian homologues. We found that 11 out of the 15 predicted transcripts were indeed localized in the glial periphery near the NMJ, suggesting that our meta-analysis holds strong predictive value, given that the probability of this happening by chance is ∼10^-5^ (Figure 3). Moreover, we found that in three cases (*Lac*, *gs2*, and *Pdi*), the localized transcripts are specifically required in glia for the correct synaptic plasticity in the adjacent motoneurons.

Knock-down of glial localized transcripts also causes morphological defects in the synapse (Figure 5). Interestingly, morphological effects were not always associated with defects in activity-dependent plasticity, and were also not always cell autonomous. These results highlight the complexity of intercellular communication necessary for proper synapse development and function. We conclude that our group of 1,700 mRNAs predicted to localize in *Drosophila* glia adjacent to the NMJ are likely to represent a rich resource of functionally relevant transcripts, which in many cases have associations with neurodegenerative, neuropsychiatric and other important nervous system diseases, further emphasizing the power of data interoperability in biological research.

It is important to consider whether the 1,700 transcripts that we predict are localized in the three glial sub-types associated with the NMJ, are in fact a core localized glial transcriptome that is functionally active in the protrusions of any glial cell type. Certainly, the glial sub-types we characterized in *Drosophila* at the NMJ are also present, as defined by specific cell type markers, in association with neurons and synapses in the central brain. Furthermore, recent studies suggest that mRNA localization in one cell type mediated by interactions with RBPs can be predictive of the same interactions and localizations in other cell types with very different morphologies and functions (Goering, Arora and Taliaferro, 2022). It has also been shown that radial glia, astrocytes, and neurons, all of which have quite distinct morphologies and functions, share a significant overlap in the types of mRNAs localized to their protrusions (D’Arcy and Silver, 2020). Perhaps cytoplasmic peripheries of any kind of cell type share many universal properties in common, such as membrane trafficking and cytoskeletal dynamics.

### 4.2 Glial protrusion transcriptomes contain a significant enrichment of disease ontology terms related to neuropathologies

Our analysis has indicated a significant enrichment of terms related to associations with nervous system diseases among the 1,700 *Drosophila* homologues with predicted glial mRNA localization (Figure 3). Moreover, this group of genes has a statistically significant overlap with the SFARI database, which contains a list of genes implicated in Autism Spectrum Disorder (ASD). There is a growing interest in the role that glial cells may play in the mechanistic causes of diverse diseases related to nervous system development and function. Indeed, it has been suggested that mRNA localization and local translation could cause glial-induced pathological effects (Sloan and Barres, 2013; Prater, Latimer and Jayadev, 2022). Our results bolster this idea and the dearth of literature in this area only serves to emphasize the need for more experimental work with a holistic approach to the nervous system, including glia and their communication with neurons at tripartite synapses. Although aggregated research made available in databases like SFARI mostly focuses on neuronal studies (Banerjee-Basu and Packer, 2010), our work shows that many SFARI genes are transcribed into mRNA that is localized to glial protrusions. We have also shown that the reactome-pathway analysis of glial localized transcripts uncovers many enriched terms related both to signaling and mRNA metabolism, providing hints at potential unexplored mechanisms related to nervous system disorders. Another highly enriched reactome pathway term worth highlighting is the “innate immune system”, which indicates that the availability of gene lists like ours is potentially valuable for multiple fields of research, as they extend beyond the context of the nervous system. Analysis modes like ours could prove useful in predicting which mRNAs might be localized in immune cells based on whether they are found in neurites.

An important direction of future research will be to characterize local translation at the peripheral cytoplasm of glia in response to signaling and neuronal activity, as was done in one recent study, which we included in our meta-analysis (Mazaré *et al*., 2020). The *Drosophila* larval system is a particularly good experimental system to study glia in relation to plasticity, because of the ease of access to the NMJ and the ability to precisely assay the role of glia in the plasticity of the motoneurons. Moreover, the extensive genetic toolbox available in *Drosophila* makes the model very tractable and well suited to studying the mechanistic effects of the introduction of disease mutations specifically into synaptic glia.

### 4.3 Vertebrate localization data meta-analysis correctly predicts mRNA localization for the majority of transcripts

Using smFISH we confirmed, with greater precision, the presence of 11 transcripts of interest in *Drosophila* glial projections, which we had discovered in a recent survey of 200 transcripts across the larval nervous system (Titlow *et al*., 2022). We have focused on the glial cell subtypes that reach the *Drosophila* NMJ because of their extensive morphologies and close contacts with the NMJ synapses (Figure S1). The NMJ associated glial subtypes are also present in the central brain, but unlike in the larval brain, in the NMJ the individual glial projections can be observed under the microscope together with individual synapses. Building on our prior work (Titlow *et al*., 2022), we confirmed that the mammalian localization data, which we re-analysed and intersected with *Drosophila* glia, correctly predicts glial mRNA localization in *Drosophila*. We conclude that peripheral glial mRNA localization is a common and conserved phenomenon and propose that it is therefore likely to be functionally important. Naturally our work is limited by the number of transcripts that could be tested experimentally. In the future, it will be important to continue to explore the localization of many more of the 1,700 *Drosophila* transcripts and test their functional requirement in glia as well as their potential implications for novel mechanisms of diseases of the nervous system.

## 5 Contributions, conflict of interest, funding details

### Author contributions

**Dalia S. Gala:** Conceptualization, Methodology, Validation, Formal analysis, Investigation, Data Curation, Writing - Original Draft, Writing - Review & Editing, Visualization. **Jeffrey Y. Lee:** Conceptualization, Methodology, Software, Validation, Formal analysis, Resources, Data Curation, Writing - Review & Editing, Visualization. **Maria Kiourlappou:** Methodology, Software, Formal analysis, Data Curation, Writing - Review & Editing. **Joshua S. Titlow:** Conceptualization, Methodology, Writing - Review & Editing, Supervision. **Rita O. Teodoro**: Methodology, Resources, Writing - Review & Editing, Supervision. **Ilan Davis:** Conceptualization, Resources, Writing - Original Draft, Writing - Review & Editing, Supervision, Project administration, Funding acquisition.

## 6 Figure legends

**Figure S1.**
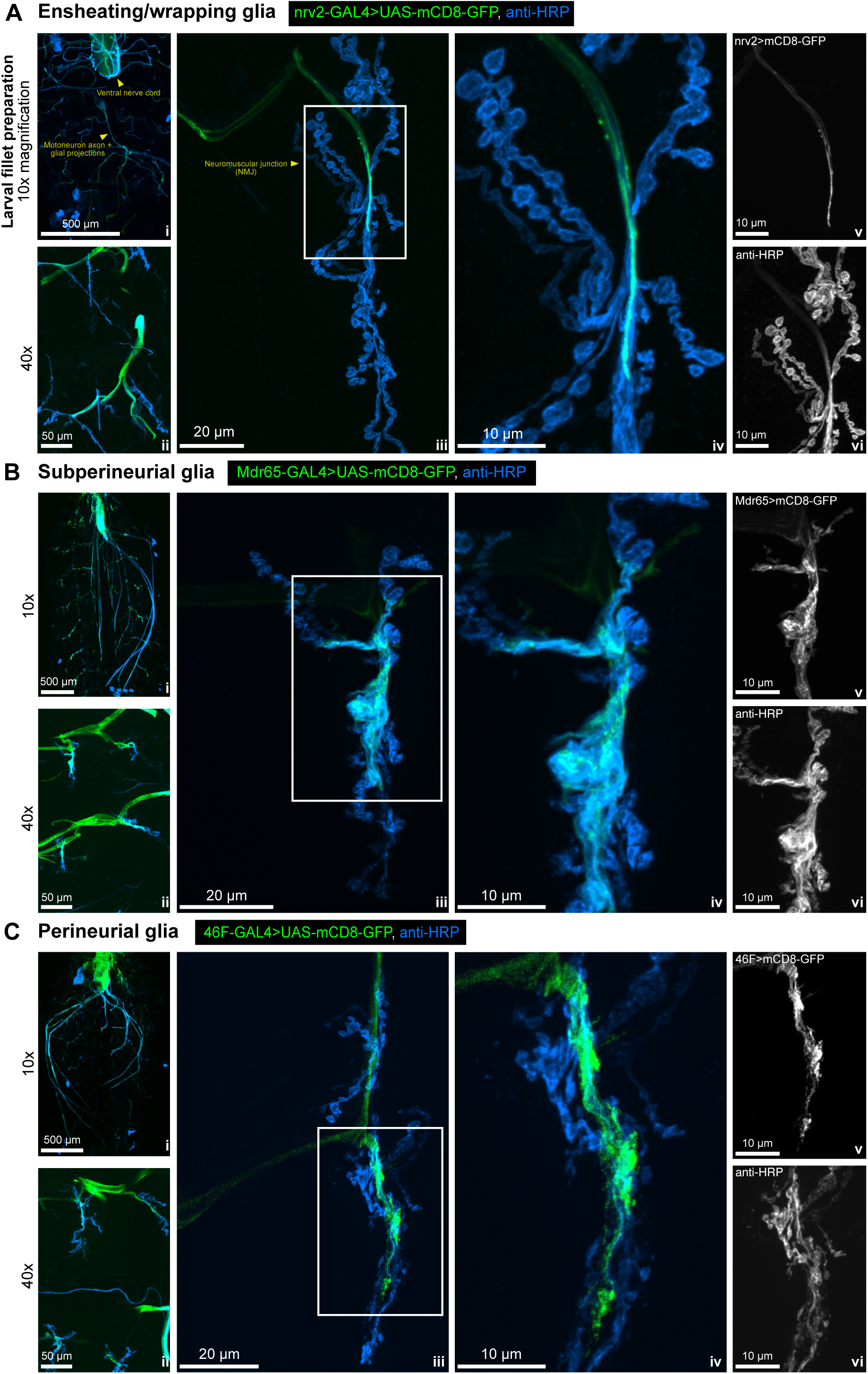
Confocal images of *Drosophila* peripheral glia. Confocal images representing the location of **A)** wrapping glia (*nrv2*-GAL4), **B)** subperineurial glia (Mdr65-GAL4) and **C)** perineurial glia (46F-GAL4) in the *Drosophila* 3^rd^ instar larva. All three of these glial subtypes produce extensive projections which reach hundreds of microns away from the cell body and reach the neuromuscular junction (NMJ) (middle two panels of each section). These three glial cell subtypes were chosen in our filtering for their elongated morphologies and the direct microscopically observable contact with the NMJ synapse. For each of A), B) and C): **i)** represents the 10x overview of the whole dissected larva, **ii)** represents a 40x zoom of segments A2 and A3 of muscles 6/7. **iii)** presents an overview of the NMJ, and **iv)** is a zoomed-in section of each of the images in **iii)**, as indicated by the white boxes. **v)** and **vi)** show the individual channels from **iv)** for clarity, with the channel label specified in the top left corner of **v)** - glia and **vi)** – anti-HRP.

**Figure S2.**
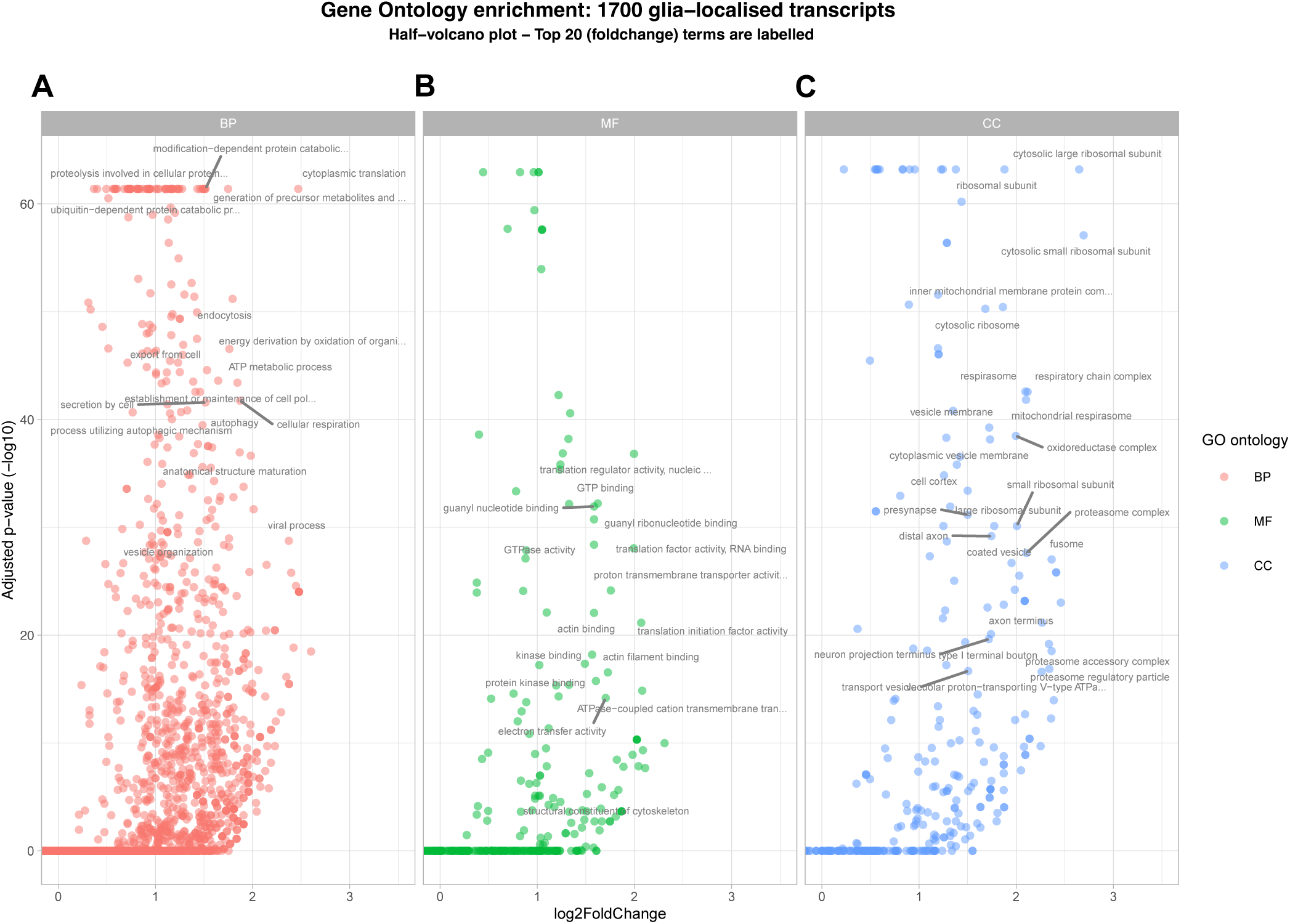
Enriched gene ontology (GO) terms of glial protrusion-localized transcripts. A-C) Volcano plot of GO terms enriched in the 1,700 glial protrusion-localized transcripts in Biological Process, Molecular Function and Cellular Component categories. *D. melanogaster* genes with high-confidence orthologues to *musculus* genes (DIOPT score ≥ 8) were used as background. Full GO enrichment analysis table is given in Supplementary Table 2.

**Figure S4.**
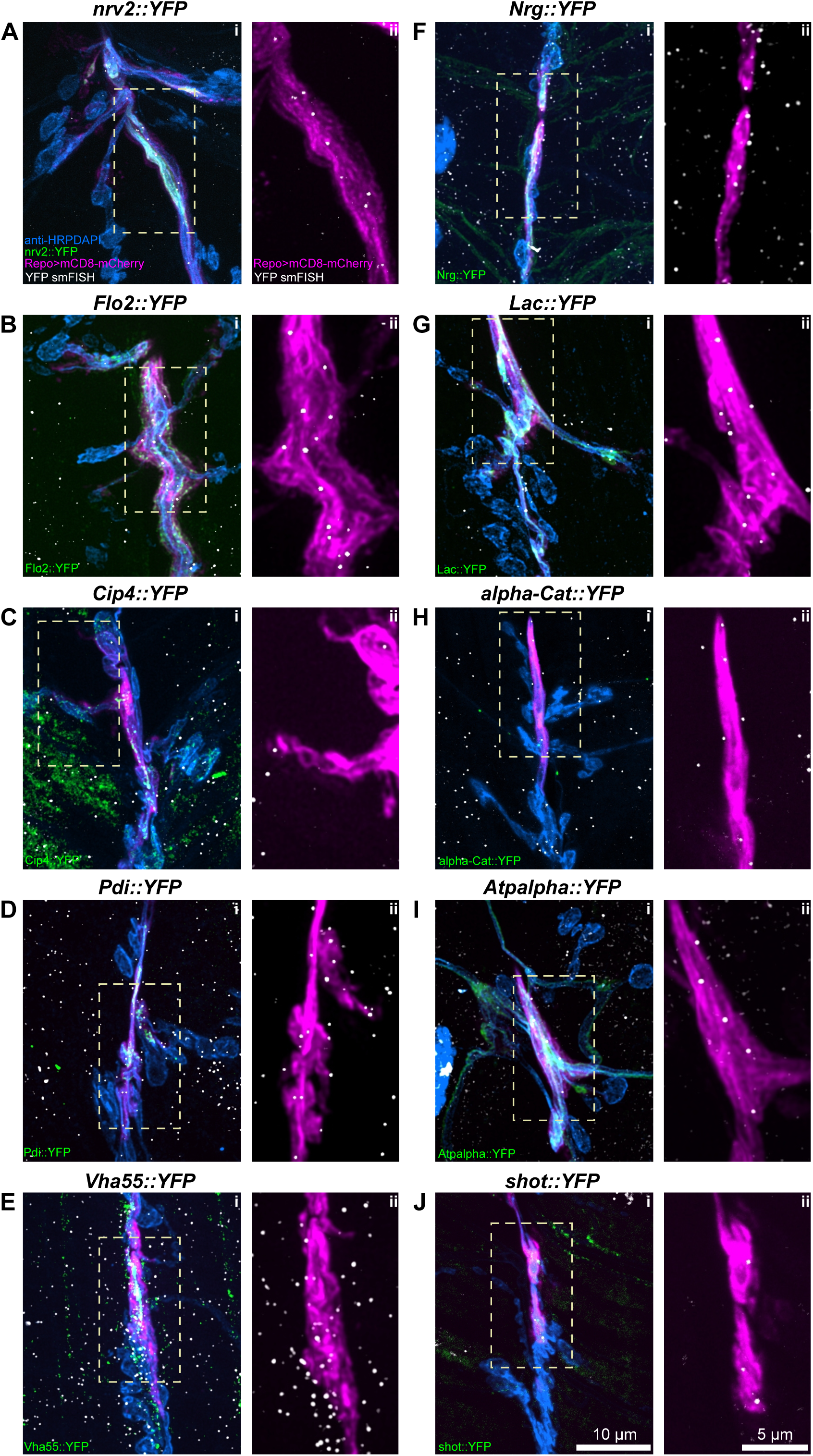
Localization of predicted transcripts in *Drosophila* glia. B-K) Confocal images of the *Drosophila* 3^rd^ instar larva NMJs (Segments A3 or A4), showing the **i)** neuron and muscle cell nuclei in blue (anti-HRP antibody conjugated to Alexa-405 fluor and DAPI, respectively), YFP conjugated proteins in green (see in-image label for details), glial membrane of perisynaptic glia labelled with Repo>mCD8-mCherry in magenta, and the YFP exon associated with the protein in white (Atto-633). For all transcripts, mRNA was detected within glial membrane labelling, as indicated in **ii)** for each panel, where a zoomed in area denoted by a white rectangle in i) is presented.

**Figure S5.**
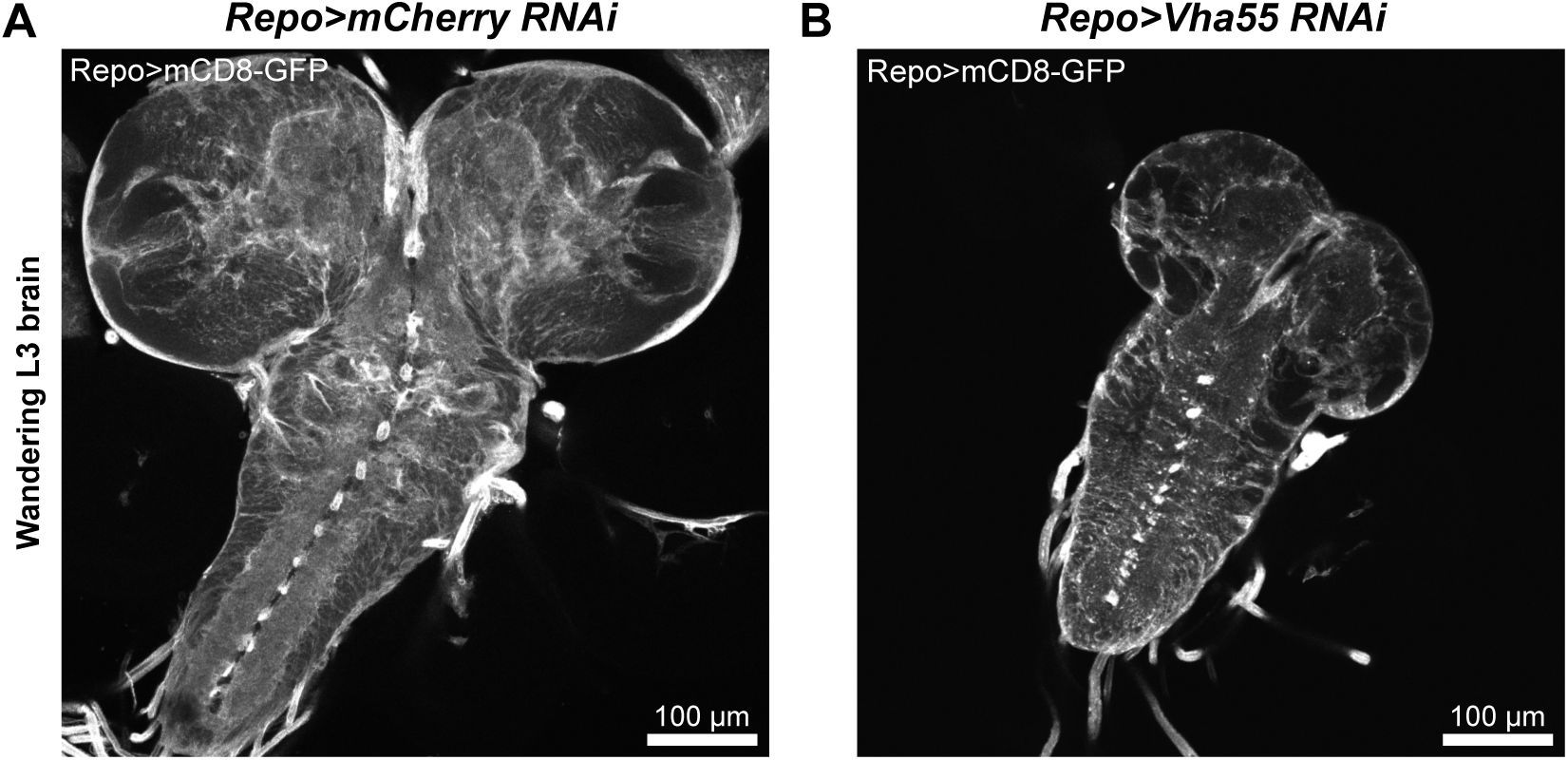
Knockdown of *Vha55* causes a small larval brain phenotype. Confocal images showing the marked difference in size between the A) control (Repo-GAL4>UAS-mCD8-GFP; UAS-*mCherry*-RNAi) and B) *Vha55*-RNAi (Repo-GAL4>UAS-mCD8-GFP;UAS-*Vha55*-RNAi) brains for 3^rd^ instar larvae of very similar sizes (representative of 5 brains for each condition).

